# Comparison of bacterial genome assembly software for MinION data

**DOI:** 10.1101/049213

**Authors:** Kim Judge, Martin Hunt, Sandra Reuter, Alan Tracey, Michael A. Quail, Julian Parkhill, Sharon J. Peacock

## Abstract

Antimicrobial resistance genes can be carried on plasmids or on mobile elements integrated into the chromosome. We sequenced a multidrug resistant Enterobacter kobei genome isolated from wastewater in the United Kingdom, but were unable to conclusively identify plasmids from the short read assembly. Our aim was to compare and contrast the accuracy and characteristics of open source software (PBcR, Canu, miniasm and SPAdes) for the assembly of bacterial genomes (including plasmids) generated by the MinION instrument. Miniasm produced an assembly in the shortest time, but Canu produced the most accurate assembly overall.  We found that MinION data alone was able to generate a contiguous and accurate assembly of an isolate with multiple plasmids.

## Introduction

The Oxford Nanopore MinION is a commercially available long read sequencer that connects to a personal computer through a USB port. Early versions of the technology showed promise for microbiological applications, including the delineation of position and structure of bacterial antibiotic resistance islands, (Ashton et al., 2014) and assembly of bacterial genomes (Loman, Quick and Simpson, 2015, Risse et al., 2015). This has been supported by the development of analysis tools for MinION data. Our objective was to compare and contrast the accuracy and characteristics of open source software for the assembly of bacterial genomes (including plasmids) generated by the MinION instrument.

Our analysis was based on a multidrug resistant *Enterobacter kobei* isolate cultured from wastewater in the United Kingdom in 2015. Genomic DNA was first sequenced using a standard protocol on the Illumina HiSeq 2000 and the short read data was assembled into ninety contigs using a Velvet-based pipeline (Zerbino and Birney, 2008). This identified a carbapenemase gene (*bla*_OXA-48_) located on a 2.5kb contig, but was insufficient to provide conclusive information on the presence and structure of plasmids. We then sequenced genomic DNA on a single MinION flow cell to determine whether long reads would allow us to resolve the structure of any plasmids present.

## Methods

### Wastewater processing and bacterial identification

Untreated wastewater was collected from a treatment plant in the UK. An *Enterobacter kobei* isolate was cultured using standard filtration and culture methods. Antimicrobial susceptibility testing was determined using the N206 card on the Vitek 2 instrument (bioMérieux, Marcy l’Étoile, France) calibrated against EUCAST breakpoints.

### Illumina sequencing and bioinformatic analyses

DNA extraction and library preparation was performed as previously described. (Quail et al., 2012) DNA libraries were sequenced using the HiSeq platform (Illumina Inc.) to generate 100 bp paired-end reads. *De novo* assemblies were generated using Velvet (Zerbino and Birney, 2008) to create several assemblies by varying the kmer size. The assembly with the best N50 was chosen and contigs smaller than 300 bases were removed. The scaffolding software SSPACE was employed (Boetzer et al., 2010) and assemblies further improved using 120 iterations of GapFiller (Boetzer and Pirovano, 2012). Species identification was based on analysis of *hsp60* and *rpoB*, as previously described (Hoffmann and Roggenkamp, 2003). To detect acquired genes encoding antimicrobial resistance, a manually curated version of the ResFinder database (compiled in 2012) (Zankari et al., 2012) was used. Assembled sequences were compared to this as described previously (Reuter et al., 2013).

### MinlON sequencing and bioinformatic analysis

DNA extraction was carried out using the QiaAMP DNA Mini kit (Qiagen, Venlo, Netherlands) following the manufacturers instructions. DNA was quantified using the Qubit fluorimeter (Life Technologies, Paisley, UK) following the manufacturer’s protocol.Sample preparation was carried out using the Genomic DNA Sequencing Kit SQK-MAP-006 (Oxford Nanopore Technologies, Oxford, UK) following the manufacturers instructions, including the optional NEBNext FFPE DNA repair step (NEB, Ipswich, USA. 6μL pre-sequencing mix was combined with 4μL Fuel Mix (Oxford Nanopore), 75μL running buffer (Oxford Nanopore) and 66μL water and added to the flow cell. The 48-hour genomic DNA sequencing script run in MinKNOW V0.50.2.15 using the 006 workflow. MetrichorV2.33.1 was used for basecalling. The flow cell was reloaded at 24 hours with the pre-sequencing mix prepared as above. MinION and Illumina sequence data have been deposited in the European Nucleotide Archive (Data citation 1)

Basecalled MinION reads were converted from FAST5 to FASTQ formats using the Python script fast52fastq.py. Read mapping was carried out to assess quality of data and coverage using the BWA-MEM algorithm of BWA v0.7.12 with the flag–x ont2d (Li, 2013). Output SAM files from BWA-MEM were converted to sorted BAM files using SAMtools v0.1.19-44428cd (Li et al., 2009).Assembly using MinION data only was undertaken using PBcR (Koren et al., 2012), Canu (Berlin et al., 2015) and miniasm (Li, 2016). Canu version 1.0 was run using the commands maxThreads=8 maxMemory=16 -useGrid=0 -nanopore-raw. PBcR pipeline with CA version 8.3rc2 was run using the options -length 500 -partitions 200 and the spec file shown in Text file 1. Minimap and miniasm were run as specified (Li, 2016). The resulting assembly was polished using Nanopolish v0.4.0 with settings as specified (Loman, Quick and Simpson, 2015), with Poretools (Loman and Quinlan, 2014) used to extract fasta sequences from fast5 files in format required by nanopolish using the option fasta. Hybrid assemblies were generated using SPAdes 3.6.0(Bankevich et al., 2012) using the option --careful, then filtered to exclude contigs of less than 1kb. All assemblies were assessed against the manually finished assembly using QUAST(Gurevich et al., 2013) version 3.2 (supplementary table 1). Assemblies were annotated using Prokka (Seemann, 2014). Figures were generated using multi_act_cartoon.py (GitHub, 2016) and MUMmer (Kurtz et al., 2004) version 3.23. Assemblies and scripts are available online (Data citation 2)

### Manually finished genome

Assemblies were generated using Canu and SPAdes as described above. A gap5 database was made using corrected MinION pass reads from the Canu pipeline and Illumina reads. Manual finishing was undertaken using gap5 (Bonfield and Whitwham, 2010) version 1.2.14 making one chromosome and nine plasmids. Icorn2 (Otto et al., 2010) was run on this for 5 iterations. The start positions of the chromosome and plasmids were fixed using circlator (Hunt et al., 2015) 1.2.0 using the command circlator fixstart. This assembly was annotated using Prokka (Seemann, 2014). The assembly and annotation is available online (Data citation 3)

## Results

Raw data was initially analysed using the Oxford Nanopore basecalling software and defined as pass or fail based on a threshold set at approximately 85% accuracy (Q9) and including only 2D reads - where data is generated from both the forward and reverse strand of DNA as it passes through the nanopore. The error rate of MinION pass data exceeded that of the Illumina data (0.048 insertions, 0.027 deletions and 0.089 substitutions per base for MinION, compared to 5.8e-06 insertions, 9.2e-06 deletions and 0.0025 substitutions for Illumina). Three tools (PBcR (Koren et al., 2012), Canu (Berlin et al., 2015), and miniasm (Li, 2016)) were used to assemble MinION pass reads alone, and a fourth (SPAdes (Bankevich et al., 2012)) was used on the combination of MinION pass data and Illumina data to produce a hybrid assembly. PBcR and Canu perform a self-correction step on reads before generating an assembly, whereas miniasm assembles the reads as provided.

All four assemblies had a similar number of contigs, and were more contiguous than the assembly using Illumina data alone, with SPAdes producing a single chromosomal contig (Table 1). We ran QUAST (Gurevich et al., 2013) to assess the quality of the assemblies, but found that it could not report all statistics for the miniasm assembly as this fell below the cut-offs for this tool. We used nanopolish (Loman, Quick and Simpson, 2015) to correct the miniasm assembly using the raw current signal (pre-basecalling) to obtain higher accuracy, and noted that the miniasm & nanopolish assembly had a similar number of indels per kilobase to Canu, although it still had more SNPs per kb (Table 1).

**Table 1.**
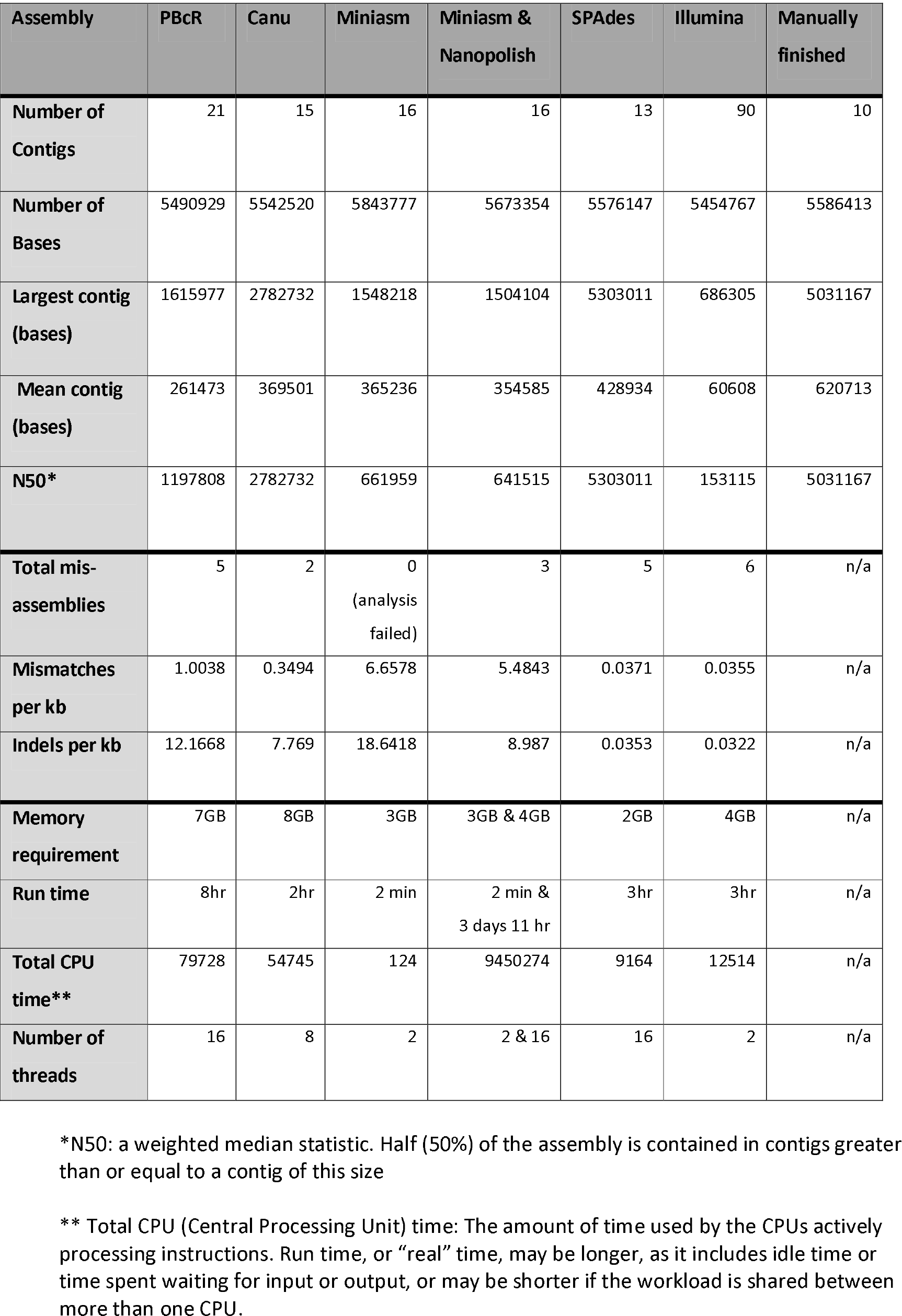
Comparison of assembly software: number and size of contigs, errors, and time/memory requirements.

| Assembly | PBcR | Canu | Miniasm | Miniasm & Nanopolish | SPAdes | Illumina | Manually finished |
| --- | --- | --- | --- | --- | --- | --- | --- |
| Number of Contigs | 21 | 15 | 16 | 16 | 13 | 90 | 10 |
| Number of Bases | 5490929 | 5542520 | 5843777 | 5673354 | 5576147 | 5454767 | 5586413 |
| Largest contig (bases) | 1615977 | 2782732 | 1548218 | 1504104 | 5303011 | 686305 | 5031167 |
| Mean contig (bases) | 261473 | 369501 | 365236 | 354585 | 428934 | 60608 | 620713 |
| N50* | 1197808 | 2782732 | 661959 | 641515 | 5303011 | 153115 | 5031167 |
| Total mis-assemblies | 5 | 2 | 0<br>(analysis failed) | 3 | 5 | 6 | n/a |
| Mismatches per kb | 1.0038 | 0.3494 | 6.6578 | 5.4843 | 0.0371 | 0.0355 | n/a |
| Indels per kb | 12.1668 | 7.769 | 18.6418 | 8.987 | 0.0353 | 0.0322 | n/a |
| Memory requirement | 7GB | 8GB | 3GB | 3GB & 4GB | 2GB | 4GB | n/a |
| Run time | 8hr | 2hr | 2 min | 2 min & 3 days 11 hr | 3hr | 3hr | n/a |
| Total CPU time** | 79728 | 54745 | 124 | 9450274 | 9164 | 12514 | n/a |
| Number of threads | 16 | 8 | 2 | 2 & 16 | 16 | 2 | n/a |

\*N50: a weighted median statistic. Half (50%) of the assembly is contained in contigs greater than or equal to a contig of this size
\*\* Total CPU (Central Processing Unit) time: The amount of time used by the CPUs actively processing instructions. Run time, or “real” time, may be longer, as it includes idle time or time spent waiting for input or output, or may be shorter if the workload is shared between more than one CPU.

Next, we compared the four assemblies to evaluate their ability to accurately reflect the genome structure. A manually finished assembly was produced and used as a reference, from which a single large inversion between the SPAdes assembly and the manually finished assembly was identified (Fig. 1). SPAdes also incorrectly integrated a plasmid into the chromosomal contig, caused by false joins. PBcR made a number of rearrangements compared to Canu (Fig. 1), validating that Canu is an improvement over its predecessor PBcR.

**Figure 1.**
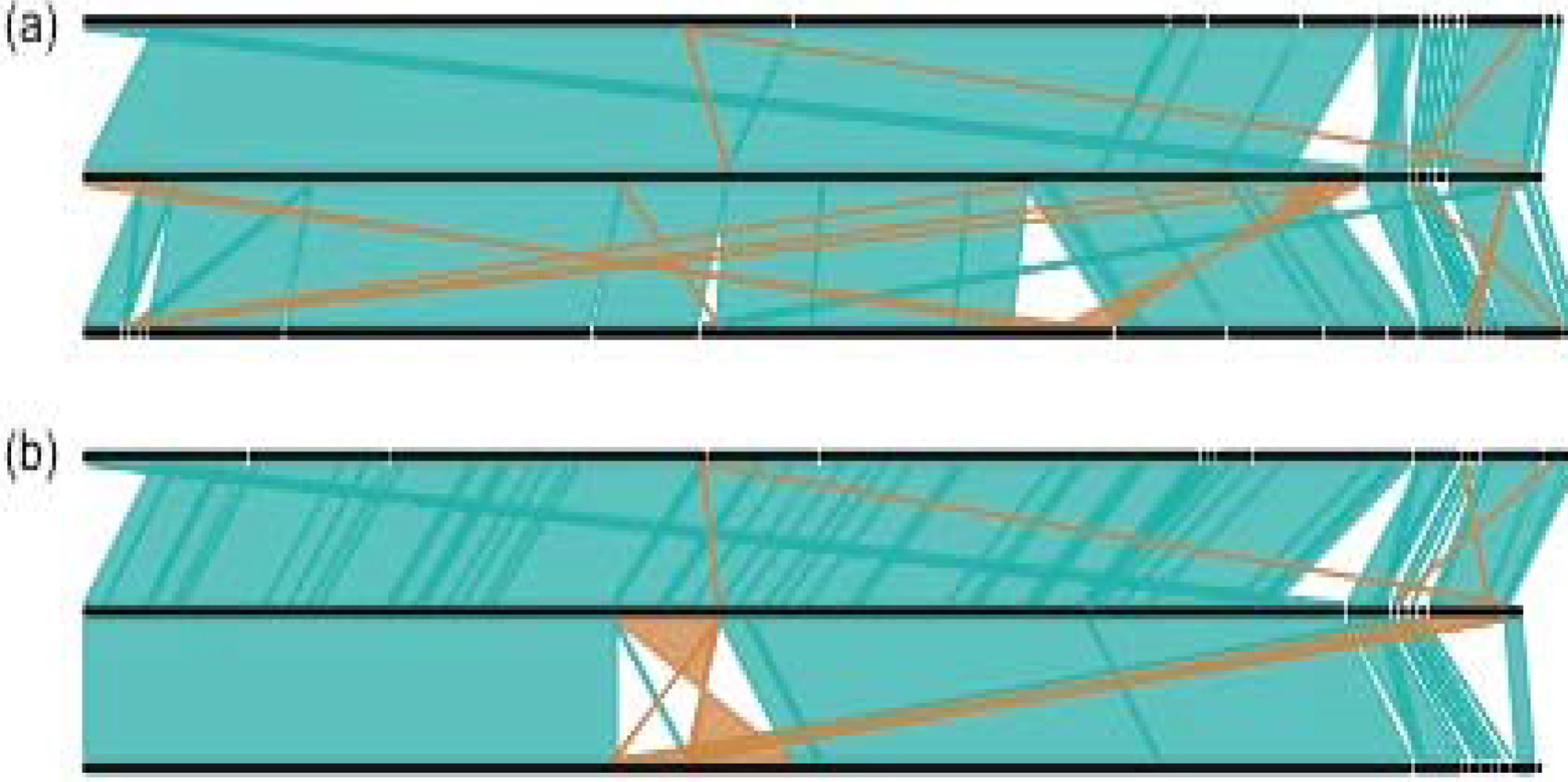
Comparison between (a) Canu, manually finished and PBcR assemblies and (b) miniasm & nanopolish, manually finished and SPAdes hybrid assemblies. Nucmer matches are shown where the length of the match is greater than lOkb or 50% of the length of the shortest sequence it matches. Forward and reverse matches are colored green and brown, respectively.

Next, we evaluated assembly of all (pass and fail) MinION reads using miniasm and Canu to determine whether adding additional (lower-quality) data would improve the assembly. Adding fail data increased the number of reads by almost 50% (64497 vs. 43260) but reduced the mean read length from 5221bp to 4687bp. Miniasm run on all reads produced the same number of contigs and a similar mean contig size as when run on pass reads. The longest contig produced with Canu was smaller when using all reads versus pass reads alone (Supplementary Table 1). With Canu, using pass reads alone led to more reads at the correction step compared to using all reads (35913 vs. 30728), indicating working with all reads could cause good quality data to be discarded during the read correction process. In both cases, using all reads did not produce a single chromosomal contig. We concluded from this that adding fail data did not consistently improve assembly.

We considered the time taken to generate sequence data, together with memory requirements to compute the assembly (Table 1). We found that almost 50% of pass reads were generated in the first 6 hours, almost 80% within 9 hours, and 90% within 12 hours. This gave a theoretical coverage of 20x, 32x and 37x, respectively. Only 31 pass reads were generated in the final 12 hours of the 48-hour run (<0.1%). Using pass reads from the first 6 hours alone led to a less accurate, fragmented assembly, but subsets of pass reads taken from the first 9 or 12 hours of the run generated comparable assemblies to pass data from the full 48-hour run (Supplementary Table 1). We also compared speed of analysis. Miniasm completed assembly within two minutes, but the trade off from using this alone was lower accuracy (Table 1). Nanopolish improved the quality of the miniasm assembly but took over three days to run; Canu took two hours and produced comparable results to the miniasm assembly after nanopolish. With current methods, isolate to assembled data in less than 48 hours is realistic. Limiting steps remaining are the requirement for overnight growth of colonies, and variable quality of flow cells.

We then evaluated the performance of the MinION to identify the presence and position of genes associated with clinically significant drug resistance in the *E. kobei* genome. HiSeq data had detected *bla*_OXA-48_ encoding carbapenem resistance on a 2.5kb contig and additional antimicrobial resistance genes in a separate 8.7kb contig (*sul1*, *arr*, *aac3* and *aac6′*-*llc*, which encode resistance to sulphonamides, rifampicin and aminoglycosides, respectively), but it was unclear whether these were on the same plasmid, two different plasmids, or chromosomally integrated. All assemblies using MinION data identified the carbapenemase *bla*_OXA-48_ on a contig with plasmid genes. The other resistance genes were identified in proximity to each other on a single large contig along with heavy metal resistance genes and plasmid genes. However, the SPAdes assembly misassembled this region into the chromosomal contig (5Mb). We concluded that there are two separate plasmids carrying resistance determinants of interest.

## Conclusion

MinION data alone was able to generate highly contiguous bacterial assemblies. Illumina data remains the cheapest way to create an assembly per sample, and still has the highest sequence accuracy, but is not without drawbacks, including the capital expenditure on the instrument or the need to outsource sequencing to a sequencing provider (which may increase turnaround time). MinION has low start-up costs, but is currently more expensive per sample. The relative ease of workflow and inexpensive laboratory set-up could facilitate its integration into routine practice, subject to improvement in reliability and reduction of MinION running cost. Whole genome sequencing is equally effective irrespective of genus, meaning that these methods could also track dissemination of plasmids containing carbapenemase genes in species such as *Pseudomonas aeruginosa*. MinION only assemblies were of sufficient quality to detect and characterise regions of antimicrobial resistance and could be generated rapidly in an outbreak.

## Acknowledgements

We gratefully acknowledge Catherine Ludden, Theo Gouliouris and the staff at the wastewater treatment plant for assistance in sample collection. We are grateful for assistance from the library construction, sequencing and core informatics teams at the Wellcome Trust Sanger Institute. We thank Simon Harris for his advice on assembling long-read data, and the staff at Oxford Nanopore for their technical support and advice during the MinION Access Program.

---

**Text 1:**
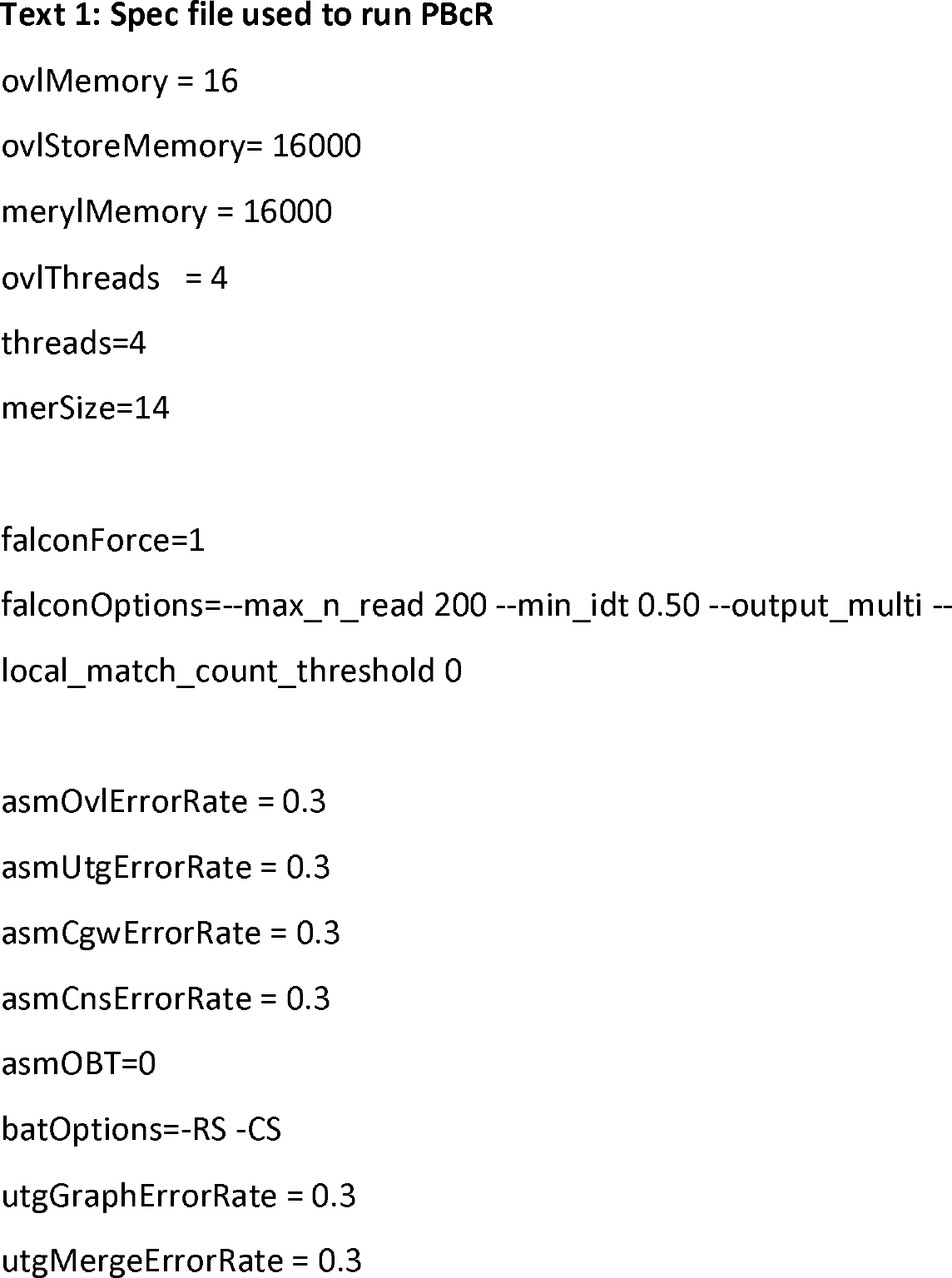
Spec file used to run PBcR.

